# Cross-Modal Scaffolding: Music Enhances Hippocampal Binding and Separation for Visual Sequential Memory

**DOI:** 10.64898/2026.02.03.703550

**Authors:** Yiren Ren, Vishwadeep Ahluwalia, Claire Arthur, Thackery Brown

**Affiliations:** School of Psychology, College of Sciences, Georgia Institute of Technology; Center for Advanced Brain Imaging, Georgia Institute of Technology/Georgia State University; School of Music, College of Design, Georgia Institute of Technology

## Abstract

The human brain continuously segments experience into meaningful episodes while also encoding temporal relationships between events, yet the mechanisms that optimize this dual challenge remain poorly understood. Here we tested a theoretical framework in which structured temporal context from one modality (music) can organize such memory computations in another (visual) through coordinated modulation of the hippocampus. Using fMRI and a sequence learning paradigm, we show that musical accompaniment enhanced both boundary detection and sequential organization of visual event memory. Mechanistically, musical context accelerated development of neural responses to boundaries in hippocampus and prefrontal cortex while simultaneously optimizing hippocampal representational patterns for learning: strengthening pattern similarity for within-sequence items while reducing computational demands for discriminating representations of different sequences. Critically, musical context created conditions where contextual similarity became a stronger predictor of memory success than before, transforming similarity from an interference signal into a beneficial learning mechanism. Multivariate analysis further revealed that musical scaffolding enhanced hippocampal encoding of the sequential position of visual stimuli, demonstrating cross-modal transfer of temporal structure to visual sequence learning. Finally, we demonstrate functional specificity across hippocampal subfields, revealing how temporal structure cues can coordinate distinct computational processes within the memory circuit. These findings establish a framework for understanding how structured context signals like music can simultaneously optimize multiple aspects of memory organization, and provide mechanistic insight for educational and clinical interventions that have leveraged cross-modal temporal enhancement to improve human cognitive function.

## Introduction

The human brain constantly receives continuous streams of information that must be organized into meaningful episodes while simultaneously extracting underlying patterns that represent how these episodes unfold.

Successful memory organization demands two complementary processes – the brain must segment continuous experience into discrete events through effective boundary detection (Kurby & Zacks, 2008; Zacks et al., 2007) and learning the precise temporal relationships between items within each event to support sequential learning and prediction (Eichenbaum, 2017; Sherman et al., 2020).

The hippocampus plays a central role in both processes. Through pattern completion, this structure can associate and bind related elements within an episode, while pattern separation allows it to distinguish between similar but distinct episodes (Leutgeb et al., 2007; Yassa & Stark, 2011). Context shapes this process, with changes in context often defining event boundaries and shaping how memories are organized (DuBrow & Davachi, 2013, 2014). For example, items sharing context are more strongly bound in memory, while context changes create natural separation points that can enhance discrimination between events (Heusser et al., 2018). The hippocampus shows exquisite sensitivity to contextual changes, implementing pattern separation when environments change to disambiguate overlapping events (Leutgeb et al., 2007) while it also encoding temporal positions within learned sequences (Hsieh et al., 2014).

However, a fundamental question remains: how are these hippocampal boundary and binding processes optimized by contextual traces to enhance learning and memory organization? While extensive research demonstrates contextual boundary effects on memory (Clewett et al., 2019; Davachi & DuBrow, 2015), we understand far less about how structured temporal context might enhance both boundary detection and sequential learning simultaneously. This gap is particularly large for statistical learning – extracting probabilistic patterns from continuous input (Saffran et al., 1996; Turk-Browne et al., 2009) – where traditional models focused primarily on pattern extraction mechanisms, with less attention to how external temporal structure might scaffold this process.

Music is a uniquely powerful example for investigating this temporal scaffolding on memory organization. Music is ubiquitous in daily life and contains inherent temporal structure at multiple hierarchical scales (Lerdahl & Jackendoff, 1983). In particular, familiar music may provide an ideal case for investigating cross-modal temporal scaffolding because of its highly temporally predictable structure – listeners can anticipate both the moment-to-moment progression and the locations of phrase boundaries that segment the musical stream. Indeed, our previous work demonstrates that familiar music enhances explicit sequence learning compared to temporally matched monotonic tones, suggesting that structured temporal predictability, not mere auditory stimulation, drives these effects (Ren et al., 2024; Ren & Brown, 2025). Music’s structured boundaries and high predictability make it an ideal stimulus for studying how structured temporal context might influence both boundary detection and pattern extraction.

Building on these insights, we used a probabilistic sequence learning paradigm to examine how musical temporal structure influences visual sequence learning. Rather than presenting deterministic sequences (as in our previous studies), we embedded probabilistic patterns in continuous visual streams, creating a more naturalistic learning scenario where participants had to extract regularities from noisy input. Some sequences were accompanied by familiar melodies, while others were presented in silence, allowing direct examination of music’s additive contribution to hippocampal memory organization. We made this design choice to fill our prior works’ gap of lacking silence, the most natural learning environment, as control.

We hypothesized that musical context would enhance sequence learning through two complementary hippocampal mechanisms (**Figure 1C**). **Mechanism 1** is **enhanced segmentation** – musical provides salient temporal boundaries that chunk visual experience into meaningful episodes. Musical phrase boundaries should trigger enhanced neural responses in hippocampus and associated regions (e.g., inferior frontal gyrus; Burunat et al., 2024; Ezzyat & Davachi, 2014; Sridharan et al., 2007), facilitate boundary detection and shape hippocampal representational geometry by strengthening association between items from difference phrases (Ezzyat & Davachi, 2014; Wallenstein et al., 1998). **Mechanism 2** is **enhanced positional encoding**. Building on evidence that the hippocampus encodes temporal sequence information (Hsieh et al., 2014; MacDonald et al., 2011) and that consistent context can enhance memory (Cox et al., 2021), we predicted that musical context would provide temporal “scaffolding” cues for encoding sequential relationships, leading to more consistent hippocampal representations of item positions within sequences (analogous to how spatial context supports place coding) and altered pattern-memory relationships that favor associative binding over interference. Using high-resolution fMRI, detailed analysis of hippocampal activity patterns, and computational modeling we tested these predictions. Critically, these two processes are complementary rather than contradictory: segmentation organizes *which* items belong together (via boundaries), while positional encoding specifies *how* items are ordered within segments (via temporal relationships).

**Figure 1:**
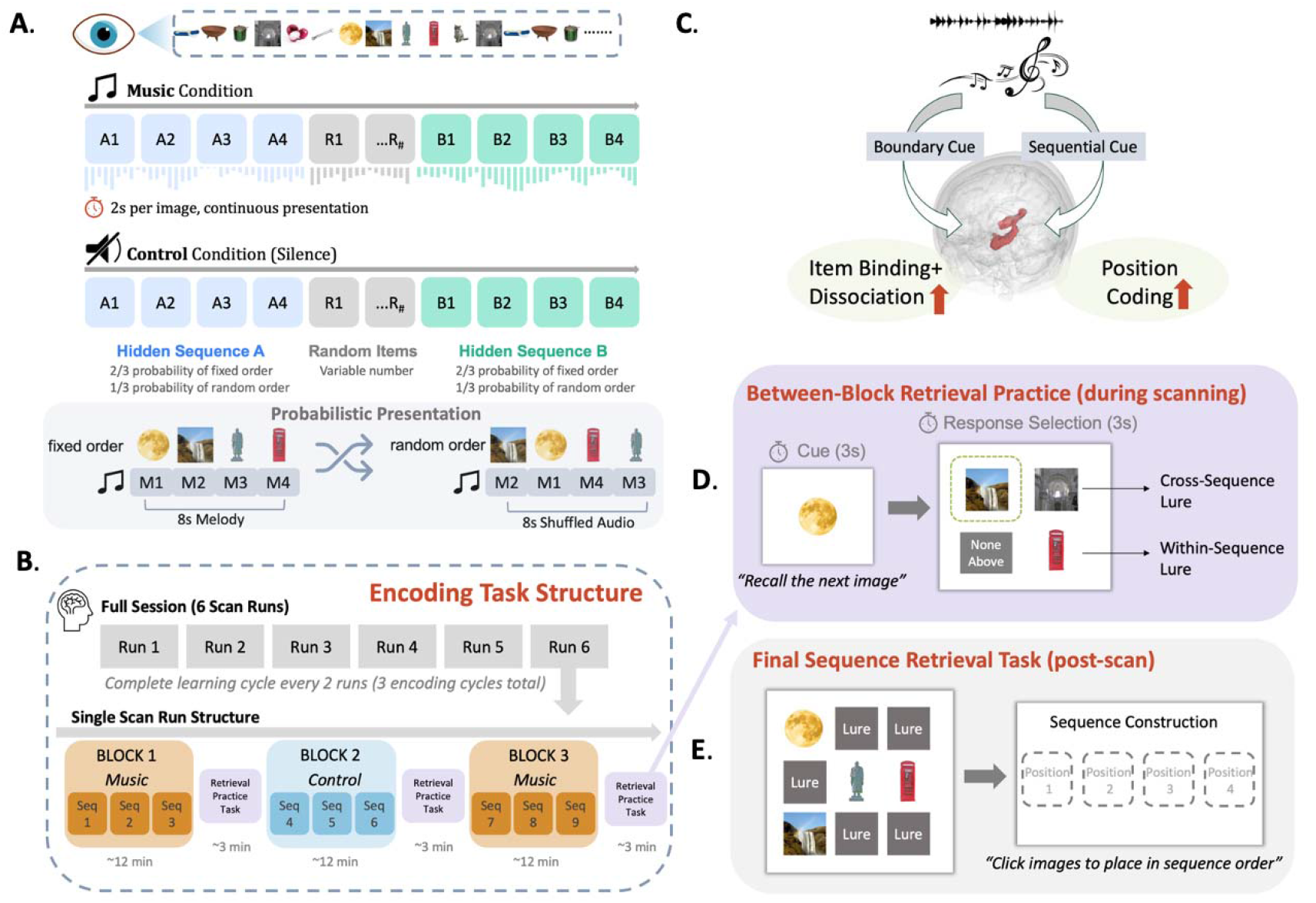
Experiment Procedure. **(A) Probabilistic Sequence Learning Paradigm**. Participants viewed continuous streams of images containing embedded 4-item sequences. In the music condition, each sequence was consistently paired with a familiar melody, with each image aligned to a specific 2-second musical phrase. In the control condition, sequences were presented in silence. Sequences appeared in correct order with 2/3 probability and in shuffled order with 1/3 probability. When sequences were shuffled, musical accompaniment was correspondingly reordered to maintain image-music pairing. Random images were interspersed between sequences to prevent simple temporal segmentation strategies. **(B) Encoding Task Structure**. The experiment consisted of six scanning runs (∼12 minutes each), with participants completing one full learning cycle of all 18 sequences every two runs (three encoding cycles total). Each run contained three blocks separated by retrieval practice tasks (**Panel D**). Within each run, two blocks used music condition and one used control condition, presented in randomized order. **(C) Theoretical Framework**. Musical context was hypothesized to provide dual benefits: boundary cues for event segmentation and sequential cues for positional encoding, leading to enhanced hippocampal item binding/dissociation and position coding. **(D) Between-Block Retrieval Practice**. After each encoding block, participants completed brief memory tests where they viewed a cue image and selected the correct next item from four options, providing online measures of sequence learning. **(E) Final Sequence Retrieval Task**. Following scanning, participants reconstructed complete sequences by selecting and ordering images from larger sets containing both target and lure items, testing subsequent sequence memory.

Our findings demonstrate that musical temporal structure enhances visual sequence learning through both predicted mechanisms. Musical context accelerated boundary detection, optimized hippocampal representational patterns for learning, and enhanced encoding of sequential position, establishing a mechanistic framework for understanding cross-modal temporal scaffolding in memory organization.

## Method

### Participants

Thirty-five healthy individuals (24 females, 12 males, mean age = 21.58 years, SD = 4.1 years, range 18-35 years) participated in the fMRI study. An additional fifty-three participants (in total 42 males and 47 females) completed behavioral testing only to validate task effects and ensure generalizability of findings across a larger sample. Sample size determination for the neuroimaging component was based on established best practices in fMRI research investigating hippocampal pattern similarity and sequence learning. Previous studies demonstrating reliable detection of hippocampal representational changes in temporal sequence learning tasks have successfully employed similar sample sizes (Schapiro et al., 2013). Our recent work using comparable methodology and sample size (N=36) successfully detected cross-modal effects on hippocampal sequence learning (Ren and Brown, 2025), confirming that this sample size is sufficient for detecting meaningful effects in cross-modal hippocampal learning paradigms.

Participant recruitment and screening followed a two-stage process. Initial screening excluded individuals with self-reported auditory impairments, visual deficits not correctable with standard lenses, fundamental music processing deficits (e.g., amusia), learning disabilities, attention disorders, or history of neurological or psychiatric conditions. Candidates who passed initial screening completed an online music familiarity assessment at least three days before the experimental session. This assessment evaluated their familiarity with twelve pre-selected musical pieces through multiple measures: immediate recognition speed, ability to continue the melody from a mid-point pause, capacity to sing along (either vocally or mentally), and overall familiarity rated on a 5-point scale. Only individuals demonstrating maximum familiarity (score of 5/5) with all musical pieces qualified for participation.

Additional screening for fMRI participants excluded contraindications for magnetic resonance imaging following the Georgia Institute of Technology/Georgia State University Center for Advanced Brain Imaging protocols. The study protocol was approved by the Georgia Institute of Technology Institutional Review Board, and all participants provided written informed consent before participation.

### Materials

#### Visual Stimuli

The experimental paradigm employed 72 distinct images selected to represent a diverse range of visual categories. Images were drawn from validated databases including the 360-color items dataset (Moreno-Martínez & Montoro, 2012), MIT indoor scenes collection (Quattoni & Torralba, 2009), and previously validated stimuli from our laboratory (e.g.Brown et al., 2020). Each image was standardized to 300x300 pixels while maintaining original aspect ratios. These 72 images were systematically organized into 18 sequences of four images each, with careful attention to balancing object, scene, and animal categories across conditions. To maintain ecological validity while preventing artificial regularities, sequences could contain varying category compositions but were constrained to include no more than one scene image per sequence. An additional set of 50 images matching the categorical distribution of sequence stimuli served as interleaved non-sequence items during encoding.

#### Musical Stimuli

Musical accompaniment consisted of twelve highly familiar melodies selected through systematic validation: “Blue Danube,” “Deck the Halls,” “Eine Kleine,” “Für Elise,” “Harry Potter Theme,” “Itsy Bitsy Spider,” “Jingle Bells,” “Ode to Joy,” “Alla Turca,” “Pop Goes the Weasel,” “Rudolph the Red-Nosed Reindeer,” and the “Wedding March”. Each piece was arranged to 8-second durations using Logic Pro software, with core melodic lines played on piano and standardized to 90 dB for melodies and 70 dB for harmonic components. Critically for neural analysis, each piece was divided into four distinct 2-second phrases that aligned precisely with visual sequence presentation, enabling examination of cross-modal temporal structure effects. During inter-sequence periods where random images were played, segments from Mozart’s Piano Sonata No. 16 provided consistent background accompaniment. Detailed familiarity validation ensuring maximum recognition (5/5 rating) for all participants is described in Ren et al., 2025.

### Experiment Procedure

The experiment comprised encoding and retrieval phases designed to examine hippocampal representational dynamics during musical context learning (**Figure 1**). Participants completed the encoding task during fMRI scanning, with behavioral procedures optimized for neuroimaging acquisition and analysis. Detailed behavioral procedures and validation are provided in Ren et al., 2025.

#### Task Design Overview

Participants viewed continuous streams of images while learning embedded sequential relationships, with some sequences accompanied by familiar music and others presented in silence (**Figure 1A**). The paradigm employed 18 four-image sequences: 12 paired consistently with specific musical pieces (music condition) and 6 presented without accompaniment (control condition). We selected silence as the control condition based on our previous work demonstrating that musical structure provides benefits beyond mere rhythmic cues when compared to monotonic tone sequences (Ren et al., 2024; Ren and Brown, 2025), allowing us to focus here on how musical context alters neural mechanisms relative to naturalistic baseline learning conditions. Each sequence appeared in its correct fixed order with 2/3 probability, while 1/3 of presentations were randomly shuffled, creating a statistical learning environment requiring extraction of probabilistic patterns from noisy input. To prevent simple temporal segmentation strategies, 1-3 random images were interspersed between sequences, with background music (Mozart Piano Sonata No. 16) played during these inter-sequence periods.

#### Practice Session

Prior to scanning, participants completed a 5-minute practice trial to familiarize themselves with the task structure. The practice consisted of four sequences presented in a format identical to the main task, with each sequence appearing six times and a 1/3 probability of random shuffling. Practice sequences were presented without musical accompaniment to establish baseline task understanding, and participants completed a brief sequence reconstruction exercise to demonstrate the learning objectives.

#### Encoding Phase Structure (fMRI Scanning)

The encoding phase consisted of six scanning runs (∼12 minutes each), with three learning blocks per run separated by brief retrieval practice tasks (**Figure 1B**). Each block contained three sequences (9 total per run), with two music condition blocks and one control condition block per run, presented in randomized order. This structure enabled examination of learning progression across three complete cycles, with participants completing one full learning cycle of all 18 sequences every two runs. Participants were instructed to 1) learn sequential relationships and 2) indicate perceived event boundaries (defined as ‘the end of an event’) via button press, providing behavioral measures of segmentation learning. Critically, participants were told that music was incidental and to focus solely on visual sequences, allowing assessment of implicit cross-modal effects on hippocampal processing.

#### Musical Context Implementation

As illustrated in **Figure 1A**, during control trials participants viewed visual sequences in silence, while during music trials each sequence was consistently paired with a specific familiar melody. Importantly, each image within a sequence was always paired with the same 2-second musical phrase, ensuring that when sequences were shuffled, the corresponding musical segments were also reordered accordingly. This design maintained the alignment between visual and musical temporal structure while preserving the probabilistic learning challenge.

#### Retrieval Testing

During scanning, after each encoding block, participants completed retrieval practice tasks (**Figure 1D**) where they viewed a sequence item and selected the correct subsequent image from four options (correct answer, within-sequence lure, cross-sequence lure, or “none above” for sequence-final items). These tasks served dual purposes: providing behavioral learning measures and creating temporal separation necessary for hemodynamic response modeling.

Following completion of the scanning session, participants exited the scanner and completed a comprehensive sequence reconstruction task (**Figure 1E**) requiring identification and correct ordering of complete four-image sequences from larger sets containing both target and lure items. This post-scan testing assessed subsequent sequence memory without the constraints of the scanning environment.

#### Neural Acquisition Considerations

Each image was presented for 2 seconds, with musical phrases precisely aligned to visual presentation timing. This design enabled time-locked analysis of cross-modal processing and examination of boundary-related neural responses. The probabilistic sequence structure and distributed learning across six runs provided sufficient trial numbers for robust representational similarity analysis while maintaining participant engagement over the extended scanning session.

### fMRI acquisition

Neuroimaging data were collected on a 3T Siemens Prisma system equipped with a 32-channel head coil at the GSU/GT Center for Advanced Brain Imaging. Complete acquisition protocols and technical specifications are detailed in Ren et al., 2025.

Anatomical reference images included T1-weighted data acquired via MPRAGE protocol (2500 ms TR, 2.22 ms TE, 8° flip angle, 0.8 mm^3^ isotropic voxels, 256 × 240 mm FOV) with GRAPPA acceleration (factor = 2). Additional T2-weighted images used 3D SPACE acquisition (3200 ms TR, 563 ms TE, 120° flip angle, 0.8 mm^3^ resolution, GRAPPA factor = 2) with matched field of view.

Task-related BOLD signals were measured using multiband echo planar imaging (EPI) with 5-fold acceleration. Functional volumes covered the whole brain (72 slices, 2.5 mm^3^ isotropic resolution) with rapid temporal sampling (750 ms TR, 32 ms TE, 52° flip angle, 220 × 220 mm FOV). Slice positioning followed the hippocampal long axis orientation.

High-resolution medial temporal lobe images were obtained using ZOOMit 2D turbo spin echo sequences with selective excitation (4560 ms TR, 51 ms TE, 130° flip angle, 2 mm slice thickness, 100 Hz/pixel bandwidth).

MRI-compatible headphones with noise cancellation enabled music delivery, combined with foam earplugs for participant comfort. Volume levels were individually calibrated while maintaining standardized relative intensities across stimuli.

### fMRI preprocessing

Data processing utilized the fMRIPrep 23.2.1 pipeline (Esteban et al., 2019) built on Nipype 1.8.6 (Gorgolewski et al., 2011). Standard processing included motion correction, slice timing adjustment, MNI space normalization, and physiological noise regression, with topup-based B0 distortion correction. Full preprocessing details, including all parameters and implementation specifics, are provided in Ren et al., 2025.

### fMRI analysis

In this paper, we focus on temporal dynamics and representational pattern analyses that specifically address hippocampal memory organization mechanisms. Additional general linear model analyses examining overall activation patterns are reported in Ren et al., 2025.

#### Finite Impulse Response Analysis

To characterize the temporal dynamics of neural responses during sequence boundary processing and examine how musical context might alter the timing of boundary-related activity, we employed finite impulse response (FIR) models. This approach investigated whether boundary-related neural patterns evolved with sequence familiarity and musical context, specifically examining: 1) potential shifts in boundary responses reflecting predictive or prolonged processing, 2) temporal dynamic differences between music and control conditions, and 3) boundary response evolution across learning stages, without assuming a specific hemodynamic response shape.

FIR models examined BOLD responses in an 8-second window following the onset of each sequence’s final image (boundary onset). The analysis used 1-second time bins to optimize temporal resolution while maintaining adequate signal-to-noise ratio. Four distinct conditions were modeled: control condition with ordered sequences, control condition with shuffled sequences, music condition with ordered sequences, and music condition with shuffled sequences. Each condition was modeled using eight FIR regressors corresponding to the eight 1-second time bins.

First-level analysis was performed for each run using FSL FEAT, providing individual subject-level analysis of temporal dynamics. Second-level analysis was conducted separately for each learning stage (runs 1-2, 3-4, 5-6), enabling examination of how boundary-related neural patterns evolved across the learning period. Statistical comparisons between conditions at each timepoint were corrected for multiple comparisons across the eight 1-second time bins using FDR correction. Two participants were excluded due to partial runs completed outside the scanner, resulting in N=33 for FIR analyses.

Based on our hypotheses regarding boundary processing, we focused our analysis on three key regions: the medial temporal lobe (MTL), the inferior frontal gyrus (IFG), and the ventromedial prefrontal cortex (vmPFC). These regions were selected based on their established roles in hierarchical sequence learning and boundary-related processing from previous literature (Ezzyat & Davachi, 2021; Schapiro et al., 2013). The IFG, parahippocampal gyrus and hippocampal masks were extracted using the Harvard-Oxford Atlas in FSL (Desikan et al., 2006). For the vmPFC, considering its broad anatomical coverage, we used 5mm spheres centered on coordinates from our laboratory’s prior studies (anterior vmPFC: MNI = [-2, 56, -10]; posterior vmPFC: MNI = [-5, 16, -11], Brown et al., 2016), which our ongoing work implicates in using schemas to support new learning.

To assess the functional significance of temporal differences between conditions, we examined correlations between BOLD responses at each 1-second time bin and participants’ segmentation performance within the same run. This analysis tested whether musical context enabled more efficient boundary processing by examining when neural activity was most strongly linked to successful boundary detection, and whether this temporal relationship differed between music and control conditions.

#### Representational Similarity Analysis

To examine whether musical context fundamentally alters hippocampal memory organization beyond boundary detection, we conducted representational similarity analysis (RSA) of hippocampal activity patterns during encoding. Based on our theoretical framework (**Figure 2.a**), we hypothesized that musical context would enhance associative binding within sequences while simultaneously improving pattern separation between sequences through distinct hippocampal mechanisms. To be specific, we predicted that musical context would provide structured temporal scaffolding that optimizes hippocampal pattern completion and separation processes (**Figure 2.a**). Specifically, within-sequence items sharing musical context should show enhanced pattern similarity compared to control items, while cross-boundary items with distinct musical contexts should show enhanced pattern separation (reduced similarity) compared to control items from overlapping blocks without clear musical boundaries.

**Figure 2:**
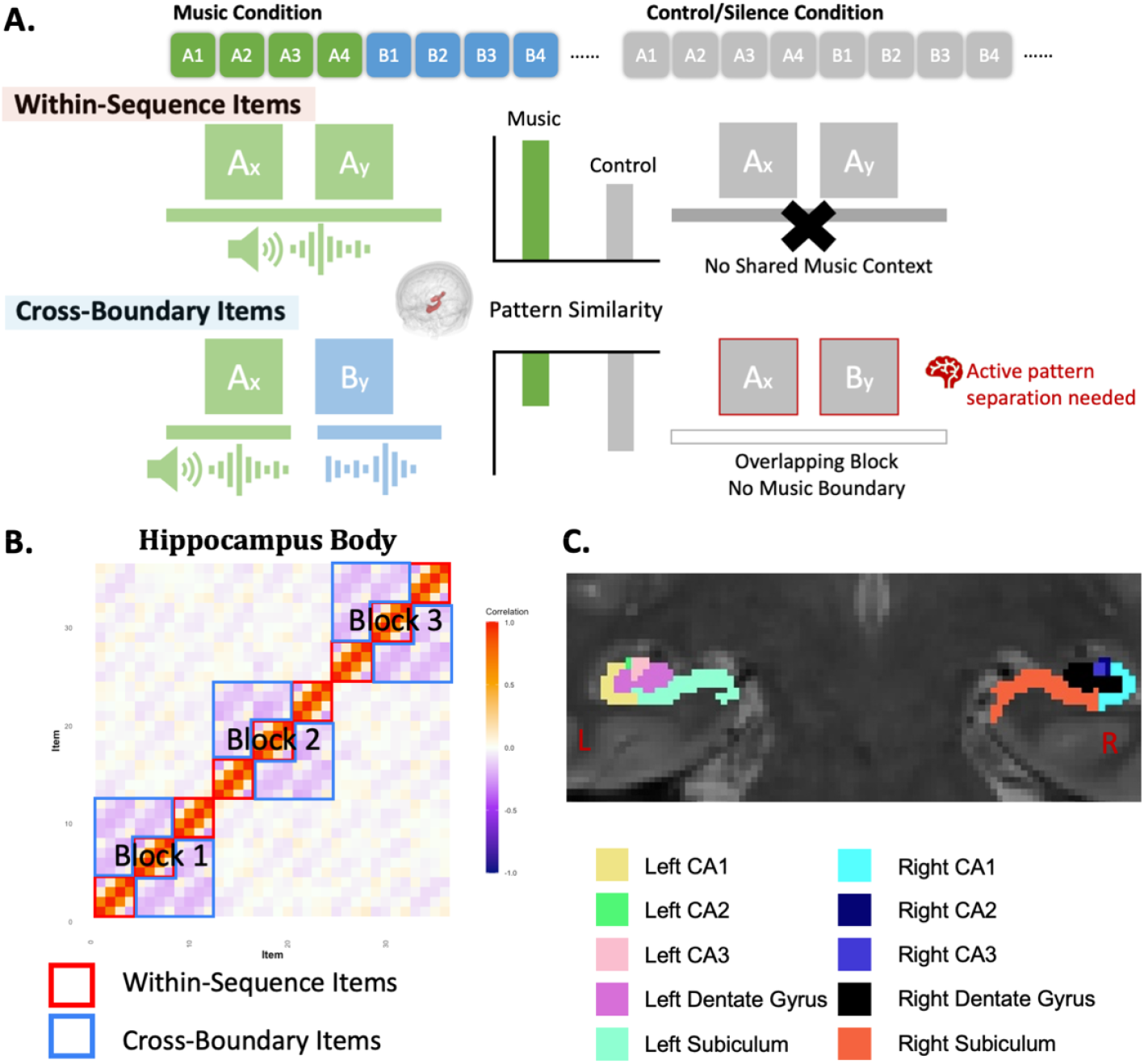
Representational Similarity Analysis (RSA) Design and Hypotheses. **(A)** Theoretical framework for how musical context modulates hippocampal pattern similarity. **Within-sequence items** (top): Items from the same sequence share musical context in the music condition, predicted to show enhanced pattern similarity compared to control items lacking shared temporal structure. **Cross-boundary items** (bottom): Items from different sequences have distinct musical contexts in the music condition, predicted to show enhanced pattern separation (reduced similarity) compared to control items from overlapping blocks without clear musical boundaries. **(B)** Representative similarity matrix from hippocampal body showing the characteristic pattern of positive correlations for within-sequence items (red boxes) and negative correlations for cross-boundary items (blue boxes) across three encoding blocks, consistent with established hippocampal binding and separation functions. **(C)** Hippocampal subfield segmentation used for detailed analysis, with each subfield color-coded to examine region-specific contributions to sequence organization and musical context effects.

To mitigate systematic biases in similarity measures, we employed separate General Linear Models (GLMs) for each of the 72 items within each run, following the “least-squares single” approach (Mumford, 2013). For each GLM, one item was modeled as the regressor of interest while remaining pictures formed another regressor, with six motion regressors included as nuisance variables. Voxel-wise beta patterns were extracted for each item from hippocampal regions of interest using unsmoothed preprocessed data to preserve fine-grained spatial patterns.

Initial analysis of the hippocampal body revealed the expected organizational structure consistent with established hippocampal binding and separation functions. **Figure 2.b** presents the averaged representational similarity matrix for 36 items in the hippocampal body (items #1-#24 were learned with music). Both music and control conditions exhibited a similar overall pattern: hippocampal patterns for items within the same sequence were positively correlated, whereas patterns for items from different sequences but learned in the same block were negatively correlated. This pattern confirms that our paradigm successfully engaged hippocampal mechanisms for associating related events while separating distinct episodes (Howard & Eichenbaum, 2015; Leutgeb et al., 2007; Rolls, 2013), providing the foundation for examining how musical context modulates these processes.

Given the distinct but complementary roles of hippocampal subfields – with dentate gyrus proposed to perform pattern separation to minimize interference between similar memories (Berron et al., 2016; Leutgeb et al., 2007), CA3 specialized for pattern completion (Rolls, 2013), CA1 for temporal sequence encoding (Barrientos & Tiznado, 2016; Hoge & Kesner, 2007), and subiculum for contextual processing such as space memory (O’Mara et al., 2009), we examined how musical context might differentially modulate these memory mechanisms across subregions (**Figure 2.c**). RSA of hippocampal subfields could reveal whether and how musical context influences both local (binding items within sequence) and global (separating items between-sequence) organizational principles of memory representations.

Hippocampal subfield masks were manually traced for each participant using ITK-SNAP (www.itksnap.org; Yushkevich et al., 2006). Our team meticulously delineated the hippocampal body, including subfields CA1, CA2, CA3, dentate gyrus, and subiculum, for both right and left hippocampus on high-resolution hippocampal scans using the finalized protocol developed by the Hippocampal Subfields Group (in preparation; preprint: Daugherty et al., 2025) (**Figure 2.c**).To align traced masks with functional imaging data, we employed a two-step registration process. First, FSL’s FLIRT was used to calculate the transformation matrix between high-resolution hippocampal scans and T1-weighted structural images, realigning traced masks into T1 native space. Subsequently, these realigned masks were transformed into MNI standard space using ANTs (Advanced Normalization Tools), ensuring spatial consistency across participants for group-level analyses. Due to motion artifacts during high-resolution hippocampal acquisition, the final sample included N=30 participants.

For each participant and run, representational similarity matrices were computed by correlating voxel-wise beta patterns across all 72 items using Pearson’s correlation within each hippocampal subregion. Items were categorized into music and control learning conditions for comparison. We focused on two critical item relationships that test our dual-mechanism hypothesis (**Figure 2.a**): (1) **Local** / **Within-sequence items** - items belonging to the same sequence, predicted to show enhanced similarity in the music condition due to shared temporal context; (2) **Global** / **Cross-boundary items** - items from different sequences learned within the same block, predicted to show altered similarity patterns in the music condition due to distinct musical boundaries providing clearer episodic demarcation.

For each hippocampal subregion, we conducted separate ANOVAs testing whether pattern similarity differed between music and control conditions for within-sequence and cross-boundary item relationships. This approach allowed us to examine how musical context specifically modulates binding versus separation processes across anatomically distinct hippocampal circuits.

To examine whether musical context altered the fundamental relationship between ordinal distance and representational similarity within sequences, we conducted gradient RSA analyses for each hippocampal subfield. For within-sequence item pairs, we coded ordinal distance as the number of positions separating items (distance = 1 for adjacent items, distance = 2 for items separated by one position, distance = 3 for items separated by two positions). We then tested whether pattern similarity varied as a function of ordinal distance using mixed-effects linear models with ordinal distance and condition (music vs. control) as fixed effects and subject as a random effect. Statistical significance was assessed using Satterthwaite’s approximation for degrees of freedom.

#### Memory-Representation Relationship Analysis

To examine how hippocampal representational patterns during encoding related to subsequent memory performance, we analyzed the relationship between neural pattern similarity and retrieval practice accuracy using the representational similarity matrices generated from the RSA analysis. For each item tested during retrieval practice, we extracted the pattern similarity value between that item and its correct subsequent item from the encoding-phase representational similarity matrices computed for the hippocampal body. This approach allowed us to examine whether stronger hippocampal pattern similarity during encoding predicted better subsequent memory for sequential relationships. We employed repeated measures ANOVAs with context condition (music vs. control), encoding run (early/middle/late), and retrieval accuracy (remembered vs. forgoteen as within-subject factors. Subject was treated as a random effect. This approach enabled examination of how musical context might alter the relationship between hippocampal representational patterns and successful memory formation across the learning period.

#### Computational Modeling and Simulation

To quantify whether our dual-mechanism hypothesis could account for the observed representational changes, we developed a Musical Scaffolding Model (MSM) that extends established frameworks for temporal context representation (Howard & Kahana, 2002) and hippocampal sequence learning (Schapiro et al., 2017). Building on Howard & Kahana’s temporal context model, we implement sequence-shared templates that create within-sequence similarity through contextual binding. Following Schapiro et al.’s approach to sequence learning, we model interference between competing sequences while adding a novel musical enhancement parameter that captures cross-modal temporal scaffolding. The MSM model (**Figure 3.a**) generates hippocampal representations by combining four components for each item *i*:

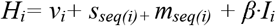

where *v*_*i*_ represents item-specific visual information, *s*_*seq(i)*_ is a sequence-shared template that creates within-sequence similarity, *m*_*seq(i)*_ is a music-specific enhancement template (active only in music condition with strength α), and *I*_*i*_ represents cross-sequence interference with strength β.

**Figure 3:**
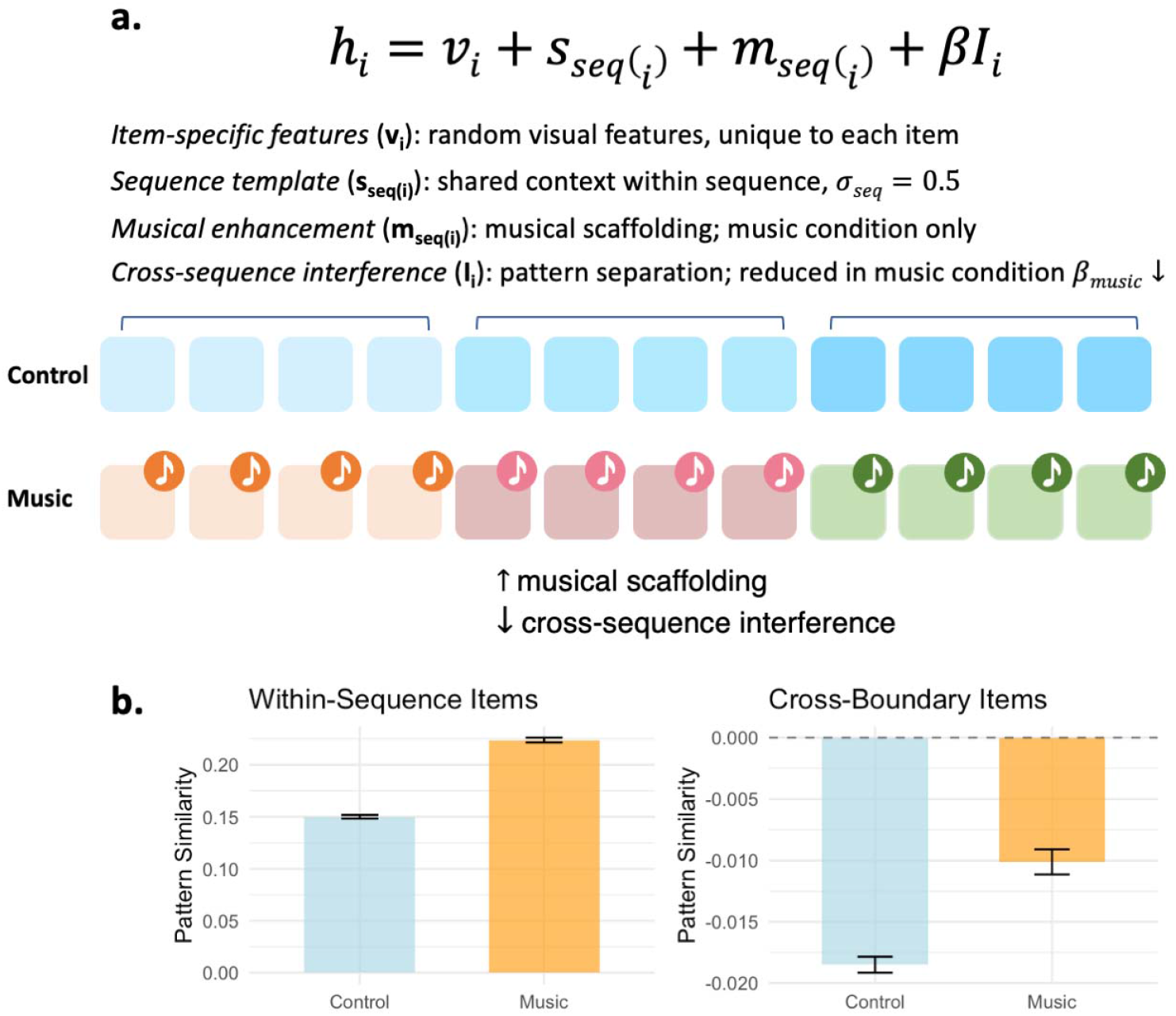
Computational model of musical temporal scaffolding. **(a)** Architecture showing how musical context enhances template sharing within sequences while reducing cross-sequence interference **(b)** Simulation results showing pattern similarity matrices or comparison with empirical data

To model within-sequence binding, each sequence was assigned a unique template *s*_*j*_∼ *N*(0, σ_seq_^2^) that was shared across all items within that sequence. This creates positive pattern similarity for within-sequence item pairs. In the music condition, sequences received additional shared templates *m*_*j*_ ∼ *N*(0, σ_mus_^2^) representing the binding effects of shared musical context. In the control condition, m_j_ = 0.

To model pattern separation between sequences, each item’s representation included interference from other sequences: *I*_*i*_ = -*β*·*∑*_*k*≠*seq(i)*_ *s*_*k*_*/(n*_*seq*_*-1*_)_, where β represents interference strength and ∑_*k*_≠_*seq(i)*_ *s*_*k*_ represents sum of sequence templates over all sequences that are different from item *i*’s sequence. This creates negative correlations between items from different sequences, with music providing clearer boundaries and thus reduced interference.

We used n_sequences_ = 6, sequence length = 4, and n_features_ = 100 to match experimental design. For each condition, we generated 100 independent simulations, computed pattern similarity matrices, and extracted within-sequence and cross-boundary correlations following identical procedures to the empirical analysis.

#### Multi-voxel Pattern Analysis

To examine whether musical context enhanced the consistency of positional encoding within sequences, we conducted multi-voxel pattern analysis using support vector machine (SVM) classification of hippocampal activity patterns. We trained binary SVM classifiers to distinguish between hippocampal activity patterns associated with adjacent sequence positions (1→2, 2→3, 3→4), testing whether musical temporal structure provided scaffolding for more distinct and consistent positional representations. We focused on binary adjacent position classification rather than multi-class position classification (1 vs 2 vs 3 vs 4) as pilot analyses revealed insufficient signal for reliable four-way discrimination. Binary classification provides greater statistical power for detecting representational differences and directly tests our hypothesis that musical context enhances sequential binding between temporally adjacent items. Classification was performed on combined left and right hippocampal body patterns to maximize signal while maintaining anatomical specificity. Classification employed leave-one-sequence-out cross-validation, where patterns from all but one sequence were used for training, and the held-out sequence was used for testing. This approach ensured that classifiers learned generalizable position representations rather than sequence-specific patterns.

Classification accuracy was compared between music and control conditions using paired t-tests, with chance performance set at 50% for binary classification. We tested whether both conditions achieved above-chance classification and whether musical context provided significant enhancement in positional discriminability.

## Results

### Musical Context Modulates Event Boundary Processing

As shown in **Figure 4.a**, musical context significantly enhanced boundary detection accuracy across all conditions. A repeated measures ANOVA revealed significant main effects of context (F(1, 1007) = 18.742, p < .001, ηp^2^ = .02), with consistently higher detection rates in the music condition for both ordered and shuffled sequence presentations. Analysis of reaction times further showed that musical context specifically accelerated boundary detection speed for ordered sequences, with participants responding 0.092 seconds faster compared to control conditions (p < .001), while shuffled sequences showed no significant timing differences between conditions (p = .493). These behavioral findings demonstrate that structured temporal context enhances both the accuracy and efficiency of event boundary detection, providing the foundation for our investigation of the underlying neural mechanisms in hippocampal memory organization.

**Figure 4:**
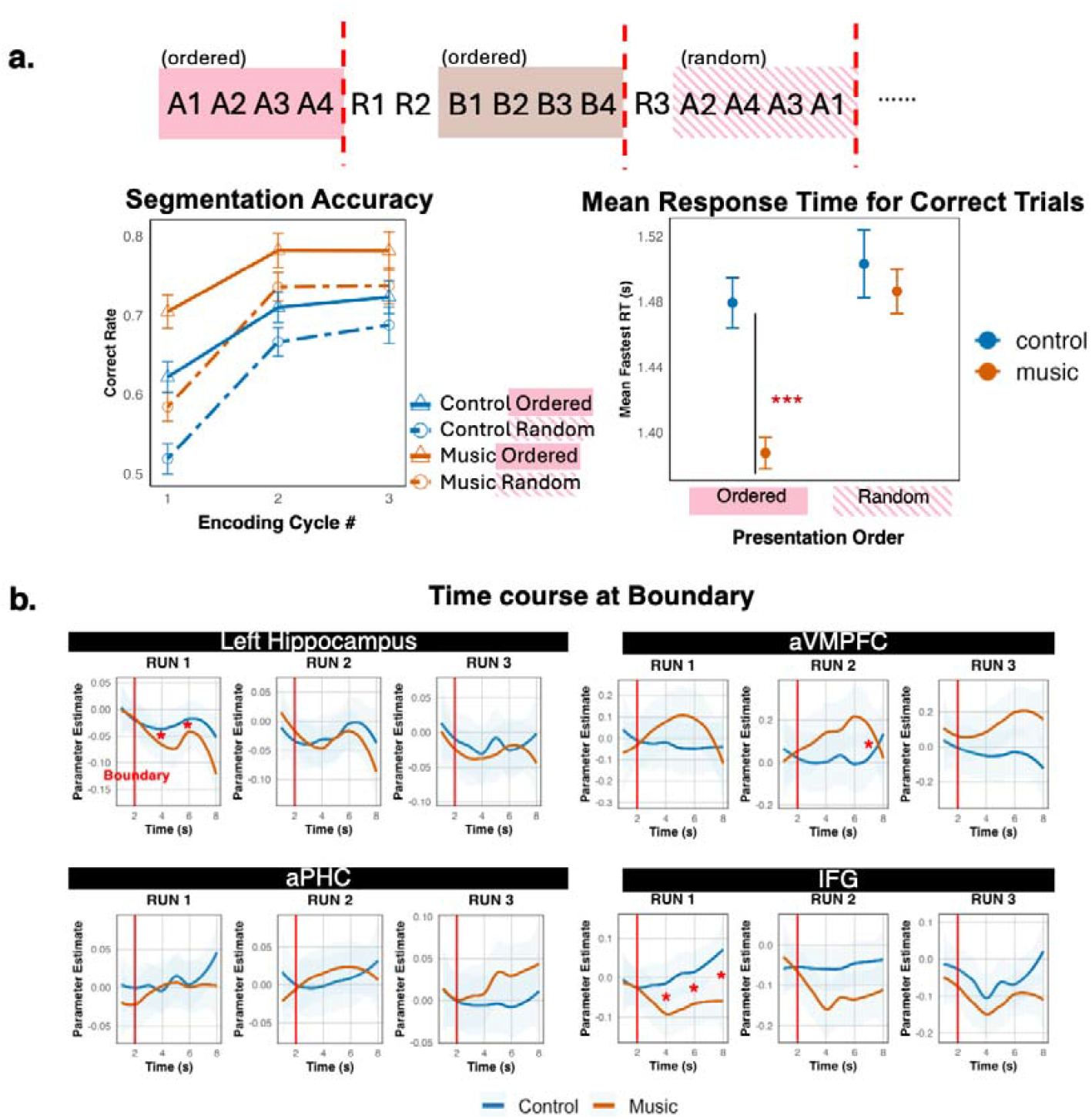
Music Enhanced Boundary Processing and Segmentation Performance. **(a)** *Segmentation task design and behavioral performance*. Participants pressed buttons to indicate perceived event boundaries in continuous image streams. Musical context enhanced segmentation accuracy in both ordered and shuffled presentation. Participants showed a higher response speed in music condition during ordered sequence presentation. **(b)** *Time Courses of BOLD Responses*: This shows the time courses of BOLD responses in the MTL, IFG and VMPFC following the onset of the last picture in each sequence. Music condition led to a stronger reduction in BOLD activity, particularly during early encoding runs, in MTL and IFG.

The time courses of BOLD responses after event boundary (for ordered presentation condition) are shown in **Figure 4.b**. we used t-tests to compare control and music conditions at each time point. In the hippocampus and inferior frontal gyrus (IFG), we observed a divergence in neural activity between music and control conditions, particularly during the early encoding run (Run 1). The music condition elicited a stronger reduction in BOLD activity shortly after boundary onset in both regions. In the hippocampus, this reduction was evident during Run 1 at Timepoint 4 (t(32) = 2.362, p = .024, pFDR = .092). This pattern suggests that music might facilitate the initial detection of event boundaries (reflected in reduced FIR response). A similar pattern was observed in the IFG, with an even more pronounced reduction at the same timepoint (t(32) = 3.251, p = .002, pFDR = .011).

The anterior ventromedial prefrontal cortex (vmPFC) demonstrated a contrasting pattern, showing significant increased BOLD activity in the music condition compared to control, particularly during the middle encoding run (Run 2). This increased activation peaked 7 seconds after the onset of the last picture (t(32) = - 3.245, p = .003, pFDR = .020).

While these findings demonstrate that musical context might enhance boundary detection, a critical question remains: do these effects reflect genuine enhancement of visual sequence learning, or merely the detection of musical boundaries themselves? To address this fundamental distinction, we examined hippocampal representational patterns during encoding to determine whether musical context systematically alters how visual sequences are organized and stored in memory.

### Musical Context Enhances Hippocampal Pattern Representations

We conducted representational similarity analysis (RSA) of hippocampal activity patterns during encoding. Given emerging evidence for distinct but complementary roles of hippocampal subfields in temporal and relational memory encoding, we examined how musical context might differentially modulate these memory mechanisms across subregions using separate repeated measures ANOVAs for each hippocampal subfield. For each subfield, we tested whether pattern similarity differed between music and control conditions for within-sequence and cross-boundary items.

#### Within-Sequence Binding Enhancement

Musical context significantly strengthened representational similarity for items within the same sequence across multiple hippocampal subfields (**Figure 5**, top panels). In right CA1, within-sequence items showed significantly higher pattern similarity in the music condition compared to control (F(1, 6234) = 4.721, p = .03). This binding enhancement was also observed in right CA2 (F(1, 5804) = 5.354, p = .021) and left subiculum (F(1, 6234) = 5.117, p = .024). In left CA3, this effect is trending to significance (F(1, 6234) = 3.81, p = .051). However, these effects did not survive FDR correction across subfields (all q > .05. See Discussion for interpretation of this modest magnitude of enhancement effects.

**Figure 5:**
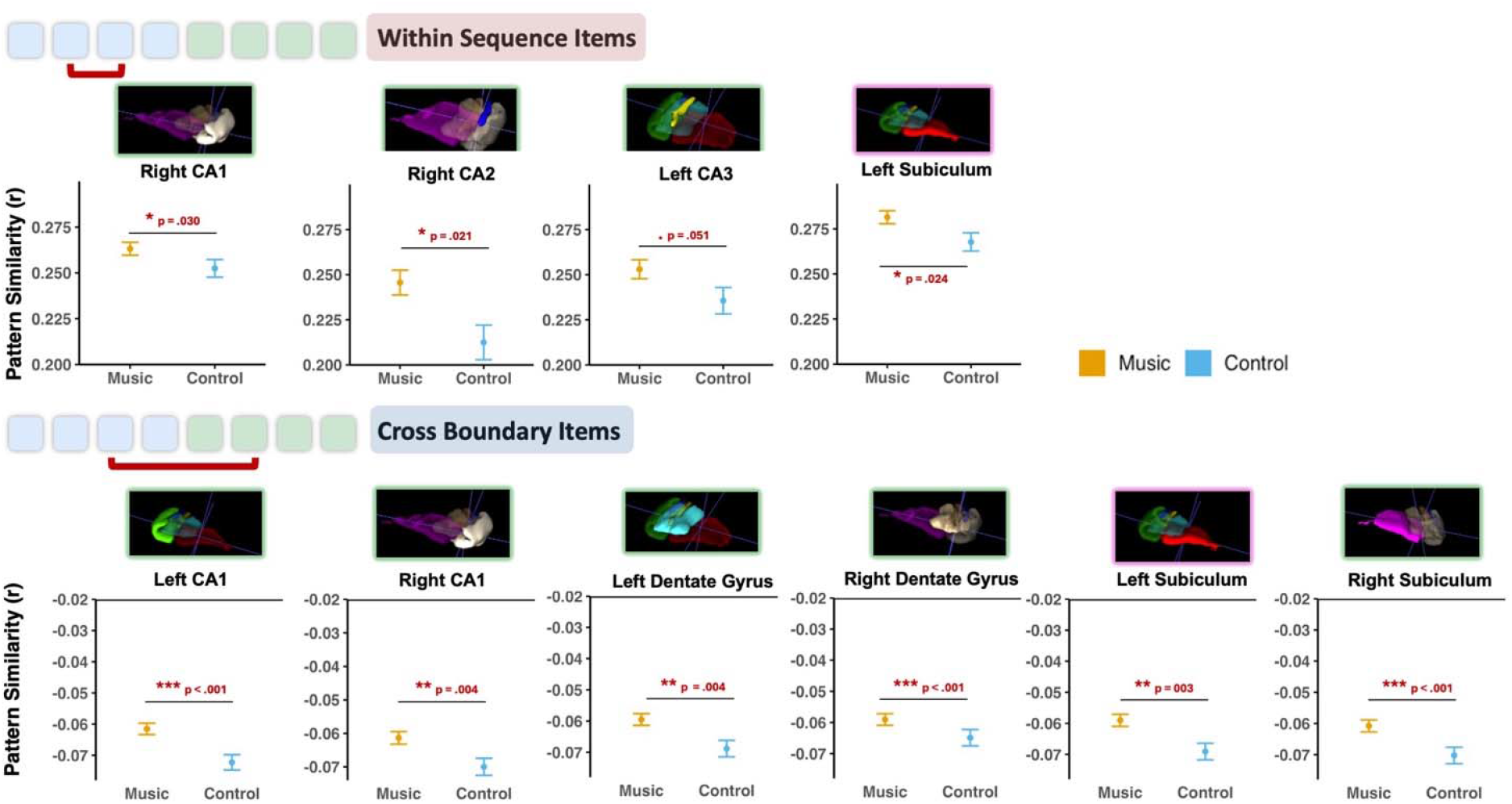
Music Enhances Both Binding and Separation in Hippocampal Subfields. **Within-sequence items** (top panels): Musical context significantly increased pattern similarity for items from the same sequence in right CA1, right CA2, left CA3, and left subiculum, indicating enhanced associative binding of temporally related events. **Cross-boundary items** (bottom panels): Musical context significantly reduced pattern similarity (increased separation) for items across sequence boundaries in bilateral CA1, bilateral dentate gyrus, and bilateral subiculum, indicating more efficient pattern separation between distinct episodes. Error bars represent ±1 SEM. Statistical comparisons: *p < 0.05, **p < 0.01, ***p < 0.001.

To examine whether musical scaffolding modulated the relationship between ordinal distance (the distance between items within the sequence) and representational similarity, we conducted gradient RSA analyses across all hippocampal subfields. Both conditions showed robust negative correlations between ordinal distance and pattern similarity (all ts < -25, ps < 2e-16), confirming position-dependent representations. Critically, music did not significantly modulate this distance-similarity gradient in any subfield (all interaction ps > .11), indicating that music did not alter the ordinal organization of items within sequence but might add a layer of temporal binding that provides modest sequence-level coherence.

#### Cross-Boundary Pattern Separation

Complementing the within-sequence binding effects, musical context altered representational patterns between sequences from different episodes (**Figure 5**, bottom panels). Cross-boundary items showed significantly reduced dissimilarity (less negative correlations) in the music condition compared to control, observed in bilateral CA1 (left: F(1, 6234) = 12.07, p < .001, pFDR < .001; right: F(1, 6234) = 8.433, p = .004, pFDR = .007), bilateral dentate gyrus (left: F(1, 6234) = 8.25, p = .004, pFDR = .007; right: F(1, 6234) = 26.29, p < .001, pFDR = .003), and bilateral subiculum (left: F(1, 6234) = 9.004, p = .003, pFDR = .007; right: F(1, 6234) = 14.07 p < .001, pFDR = .008).

#### Computational Model Validation

To quantify whether our proposed mechanisms could account for the observed hippocampal representational changes, we developed a Musical Scaffolding Model (MSM) (**Figure 3a**) that simulates musical scaffolding through enhanced template sharing and reduced interference.

Through systematic model fitting (parameter optimization) we showed that the MSM model can match empirical cross-boundary correlation patterns. To explain our results, the optimal model parameters were: σ_seq_ = (sequence template strength), σ_mus_ = 0.4 (musical enhancement strength), β_control_ = 0.35 and β_music_ = 0.2 (interference strengths). The MSM model successfully reproduced the empirical hippocampal representational patterns (**Figure 3.b**). Mechanistically, the model reveals that musical context operates through coordinated modulation of hippocampal computational processes: the enhanced template sharing parameter (σ_mus_ = 0.4) demonstrates how shared musical context strengthens representational binding within sequences, while the 43% reduction (0.35-0.2)/0.35 = 0.43)) in interference strength from the control condition shows how musical boundaries reduce the computational effort required for pattern separation between distinct episodes.

### Musical Context Transforms Neural Pattern-Memory Relationships

The retrieval practice tests inserted during encoding allowed us to examine how musical context influences both behavioral learning and underlying neural representations. We first assessed whether musical context enhanced sequence learning performance, then examined how hippocampal pattern similarity related to successful memory formation.

#### Behavioral Learning Enhancement

Musical context during encoding consistently improved next-item retrieval accuracy during between-block retrieval task **(Figure 1.d, Figure 6.a**), particularly in later encoding runs (**Figure 6.c**). A repeated measures ANOVA revealed significant effects of both run (F(2, 517) = 42.05, p < .001, ηp^2^ = .14) and context condition (F(1, 517) = 9.522, p = .002, ηp^2^ = .02). Post-hoc comparisons showed significantly higher retrieval accuracy in the music condition during run 2 (mean difference = 0.071, p = .026) and a trending advantage in run 3 (mean difference = 0.053, p = .099).

**Figure 6:**
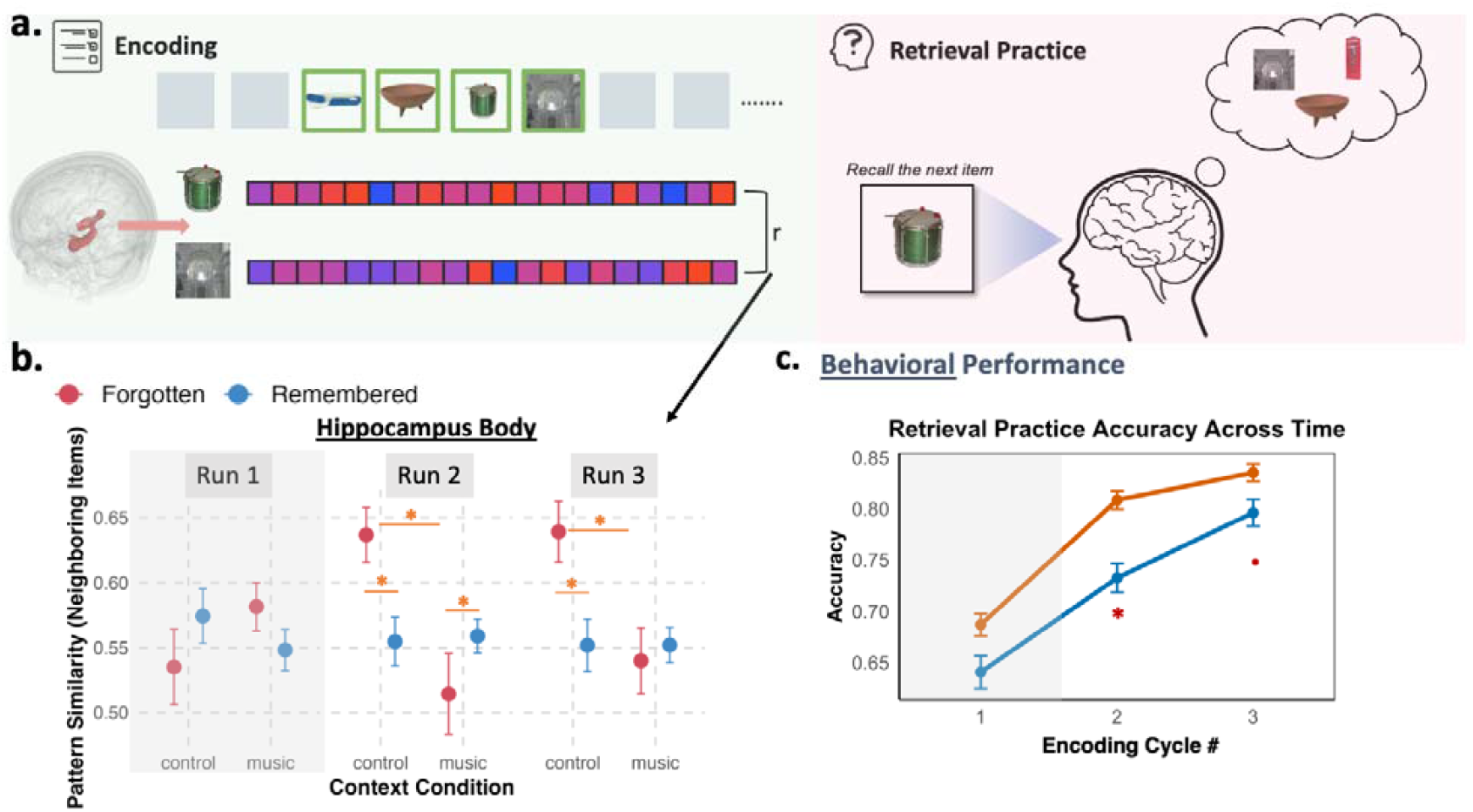
Adjacent Items Pattern Similarity and Retrieval Practice Accuracy. **(a)** Hippocampus pattern similarity during encoding for adjacent items that were subsequently tested during the same-block retrieval practice was compared for correctly remembered items versus forgotten items **(b)** Hippocampal pattern similarity for adjacent items during encoding, separated by subsequent memory performance. Musical context led to higher pattern similarity for item pairs that were later remembered during retrieval practice, in runs 2-3, while the control condition showed the opposite pattern. **(c)** Retrieval practice accuracy across encoding cycles, demonstrating progressive learning in both conditions. However, musical context condition showed consistently higher retrieval practice accuracy especially in run 2-3.

#### Neural Pattern-Memory Relationships

To examine how hippocampal representations related to successful learning, we analyzed pattern similarity between adjacent items during encoding as a function of subsequent retrieval performance. In the hippocampus body, pattern similarity was significantly modulated by the interaction of context and run (F(2,2404.7) = 5.576, p = .004) and by the three-way interaction of context, run, and accuracy (F(2,2408.3) = 5.686, p = .003). Critically, during runs 2-3, the music and control conditions showed opposite patterns (**Figure 6a-b**). Post-hoc comparisons revealed that, in the control condition, higher hippocampal pattern similarity between adjacent items was associated with subsequent forgetting (run 2: estimate = .079, t(2400) = 2.018, 0 =.044; run 3: estimate = .086, t(2397) = 2.151, p = .032), whereas in the music condition during run 2, higher similarity associated with successful remembering (trending: estimate = -.0.468, t(2408) = -1.842, p = .065). Forgotten trials adjacent items pattern similarity is significantly higher in the control condition than music condition (run 2: estimate = .0.1223, t(2408) = 2.818, 0 =.005; run 3: estimate = .1, t(2404) = 2.261, p = .024).

### Musical Context Provides Sequential Scaffolding

Having established that musical context optimizes hippocampal representations during encoding, we examined whether these effects translate to enhanced sequential memory and more consistent positional encoding in the hippocampus.

#### Enhanced Final Sequence Retrieval

Following the encoding session, participants completed a comprehensive sequence reconstruction task (**Figure 7.a**). Musical context significantly enhanced sequence reconstruction performance. Participants successfully reconstructed 57.8% of music-accompanied sequences (SD = 30.6%) compared to 52.5% of control sequences (SD = 31.7%), representing a meaningful improvement in sequential memory accuracy (t(86) = 2.292, p = .024, d = .246). This enhancement demonstrates that structured temporal context during encoding strengthens the formation of sequential associations that persist beyond the learning session, confirming that musical scaffolding produces lasting benefits for temporal order memory.

**Figure 7:**
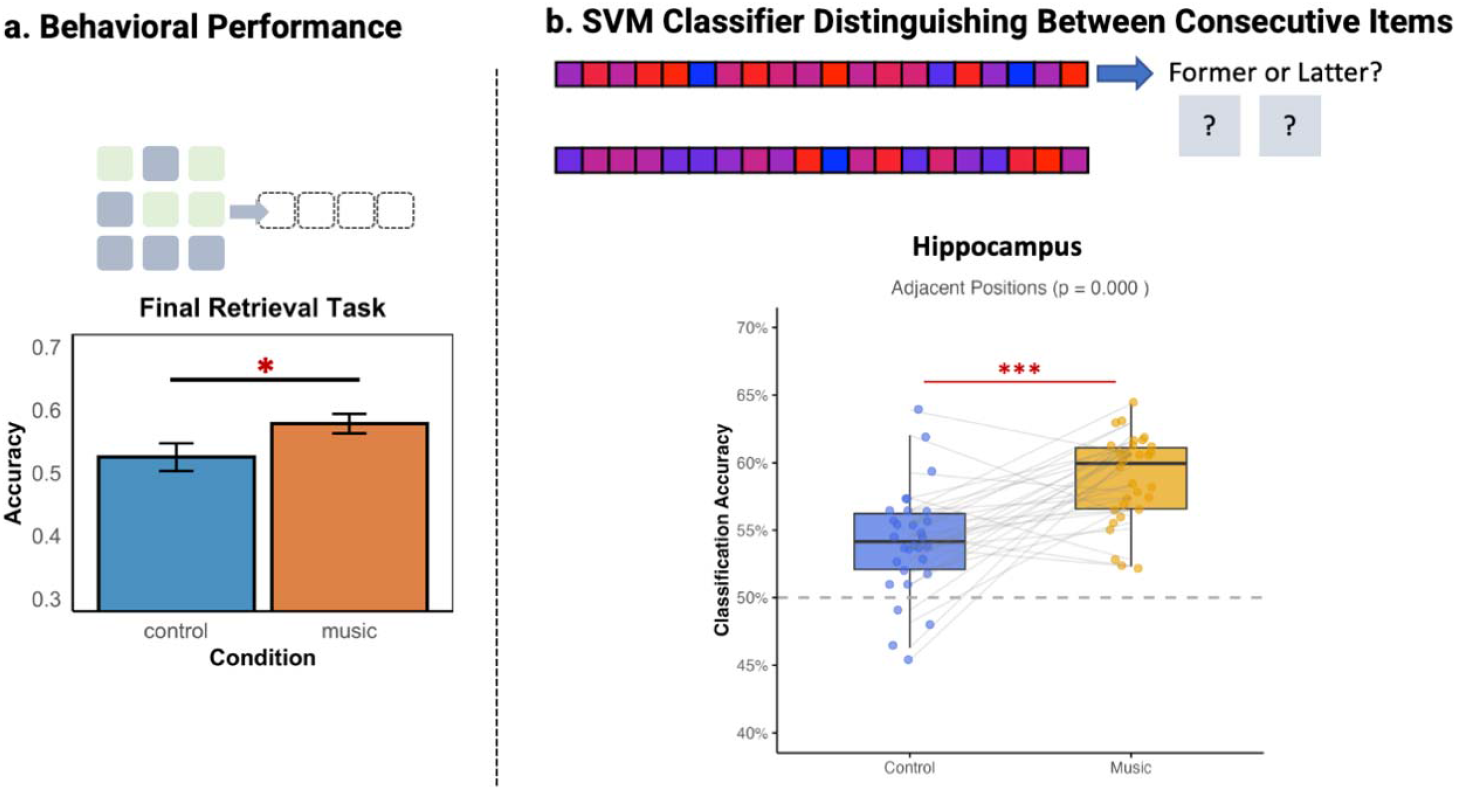
Sequential Position Scaffolding and Memory Performance. **(a)** Final sequence retrieval performance after scanning. Participants reconstructed complete 4-item sequences by selecting and ordering images from a larger set. Musical context led to significantly better sequence reconstruction accuracy. **(b)** Multivariate classification of sequential positions using hippocampal activity patterns. Support vector machine classifiers trained to distinguish between consecutive positions within sequences showed significantly higher accuracy in the music condition, indicating more distinct and consistent positional representations. Individual data points show subject-level performance with connecting lines. Dashed line indicates chance performance (50%).

#### Improved Positional Representation Consistency

To test whether musical context enhanced the consistency of positional encoding, we used binary SVM classifiers to distinguish between hippocampal activity patterns for adjacent sequence positions. Classification accuracy was significantly above chance (50%) for both conditions (music: t(89) = 11.34, p < .001; control: t(89) = 5.626, p < .001), confirming that hippocampal patterns contain reliable positional information. Critically, the music condition showed significantly higher classification accuracy (M = 60.84%, SD = 7.4%) compared to the control condition (M = 54.1%, SD = 7%; t(89) = 4.286, p < .001, d = .452) (**Figure 7.b**).

We attempted to examine whether positional decoding effects differed across hippocampal subfields, given theoretical predictions that CA1 might show stronger position coding as well as DG/CA3, supporting pattern separation. However, classification performance in subfields did not exceed chance levels, likely due to insufficient voxel counts or limited resolution.

## Discussion

### Overall Summary

This study demonstrates mechanisms behind a fundamental principle of cross-modal temporal scaffolding: how familiar temporal context from one modality can systematically optimize memory organization in another through coordinated hippocampal mechanisms. Comparing familiar music to silence revealed dual complementary processes (**Figure 1C**): enhanced segmentation that strengthened within-sequence associations while improving between-sequence discrimination, and enhanced positional encoding that provided organizational frameworks for temporal relationships. These findings demonstrate that temporally predictable auditory context can scaffold visual sequence learning through hippocampal mechanisms.

### First-Level Facilitation: Enhanced Boundary Processing

Musical context altered the temporal dynamics of boundary processing across memory and control networks. FIR analysis revealed earlier and stronger BOLD signal reductions in hippocampus and inferior frontal gyrus right after the onset of boundary image, while anterior vmPFC showed delayed activation enhancement after boundary. This temporal reorganization likely reflects enhanced boundary detection facilitated by the predictable temporal framework provided by familiar melodies. Musical phrase endings coincided with visual sequence boundaries, creating salient transitional signal (Burunat et al., 2024; Koelsch et al., 2019; Zacks et al., 2011) that were temporally aligned with the visual modality and may have strengthened boundary representations.

The anatomical specificity of these effects provides additional mechanistic insight. The hippocampus showed enhanced boundary-related activity in the music condition, aligning with its role in boundary detection and encoding of temporal relationships (Ezzyat & Davachi, 2011). The concurrent involvement of IFG aligns with its established role in boundary detection (Sridharan et al., 2007) and multisensory binding during rule learning (Li et al., 2020), suggesting coordinated engagement of memory and prediction systems when temporal structure is available across modalities.

These boundary detection findings provide initial evidence for enhanced segmentation, but they represent only one aspect of how musical context might optimize the brain’s ability to categorize and organize sequential information. Enhanced segmentation should also manifest at the level of hippocampal representational patterns—specifically, in how the brain encodes relationships within sequences versus between different sequences. We therefore examined whether musical temporal structure produces coordinated changes in hippocampal pattern similarity that reflect this enhanced segmentation at the representational level.

### Coordinated Optimization of Binding and Separation Mechanisms

Musical context markedly altered hippocampal memory organization through two representational mechanisms. Our RSA results revealed that musical accompaniment produced the coordinated effects predicted in our theoretical framework (**Figure 2a**): enhanced within-sequence binding coupled with altered cross-boundary separation patterns. These effects demonstrate that temporal predictability provided by familiar music can modulate hippocampal representational geometry.

<u>Enhanced within-sequence binding</u> was primarily mediated by the associative memory circuit, with robust effects in CA1, CA2, and subiculum, plus a trending effect in CA3 (**Figure 5** top panel). This pattern aligns with established roles of these regions in forming and maintaining associative representations.

CA1’s robust enhancement reflects its role in integrating temporal and contextual information (Eichenbaum, 2014), which may be enhanced by music’s additional temporal context for integration that enhances CA1’s capacity to bind items sharing the same temporal framework. CA2’s increased pattern similarity suggests enhanced temporal coding, consistent with its emerging role in sequential processing and signal transmission to CA1 (MacDonald & Tonegawa, 2021). The subiculum serves a critical role in integrating hippocampal outputs with distributed cortical regions involved in contextual encoding (Kim et al., 2012). The enhanced pattern similarity for within-sequence items observed in the subiculum during music condition likely reflects its role in strengthening contextual associations at multiple levels: both between sequential items and between these items and their broader temporal context provided by music. This interpretation aligns with recent evidence demonstrating the subiculum’s involvement in contextual binding beyond its simple relay functions (Sun et al., 2019). Lastly, while CA3 showed a trending rather than significant effect, this pattern aligns with its specialized role in pattern completion and associative memory formation (Rolls, 2013). CA3’s extensive recurrent connections enable completion of partial patterns and binding of contextually related elements (Neunuebel & Knierim, 2014; Rolls, 2013), suggesting that musical context may provide additional associative cues that facilitate these completion processes, though to a lesser degree than other subfields.

<u>Enhanced cross-boundary separation</u> was primarily driven by the pattern separation circuit, with bilateral effects in dentate gyrus, CA1, and subiculum (**Figure 5**, bottom panel). The observed decreased repulsion (elimination of dissociation) across these regions during the music condition reflects more efficient pattern separation processes. When input patterns share features and have overlapping neural representations, like in our control condition (sharing learning block and context), pattern separation can manifest as active “repulsion” between these representations—pushing them further apart in representational space (Wanjia et al., 2021). Musical context reduced this computational burden by providing clear temporal boundaries for episode discrimination. Rather than requiring maximal representational orthogonalization, the hippocampus leveraged musical structure for effortless separation—achieving better discrimination with reduced computational effort.

The bilateral dentate gyrus effects align with its established function in pattern separation and context change detection (Leutgeb et al., 2007), where orthogonalized patterns enable better discrimination between similar stimuli (Berron et al., 2016). Similarly, CA1’s bilateral involvement reflects its role in actively detecting mismatches between current input and stored representations (Duncan et al., 2012). The reduced negative cross-boundary pattern correlations in the music condition suggest that musical boundaries provide such clear contextual shifts that less active mismatch detection is required, enabling more efficient episodic discrimination. The subiculum’s bilateral effects extend its contextual integration role (Kim et al., 2012) to cross-boundary processing, likely reflecting its capacity to use musical contextual information to organize distinct episodes within a broader temporal framework without excessive representational competition.

This anatomical dissociation reveals that musical context doesn’t simply boost hippocampal function uniformly, but rather encourages different subregions to optimize and express specialized contributions to memory organization. The coordination between associative regions strengthening within-episode binding and separation regions optimizing between-episode discrimination demonstrates how external temporal structure can simultaneously enhance both aspects of segmentation through targeted recruitment of appropriate neural circuits.

The magnitude of within-sequence enhancement effects is statistically modest compared to cross-boundary effects, which may reflect the hippocampus’s fundamental role in associative binding (Staresina & Davachi, 2009). Even in the control condition, hippocampal circuits may already approach optimal binding capacity for temporally contiguous items, leaving limited room for further enhancement through musical scaffolding. This interpretation suggests that musical context provides incremental rather than transformative enhancement of an already efficient binding process.

Notably, the strong pattern correspondence between our computational model and empirical data provides mechanistic insight into how temporal scaffolding optimizes hippocampal memory organization. The MSM model demonstrates that musical context effects can be understood through two complementary parameters: enhanced template sharing within sequences and reduced interference between sequences (β reduction of 43%). This quantitative framework suggests that familiar music temporal structure provides computational scaffolding that simultaneously optimizes both pattern completion and pattern separation processes – this computational architecture reveals how external temporal structure can simultaneously strengthen ‘what goes together’ while clarifying ‘what stays apart’ in hippocampal memory organization. Furthermore, the parameterization provides testable predictions for other forms of structured context: any temporally predictable stimulus should enhance memory organization in proportion to its template sharing strength and boundary clarity, offering a quantitative framework for designing optimal learning environments.

### Cross-Modal Transfer of Temporal Organization

Having established that musical context optimizes hippocampal organization of which items belong together versus apart, we next examined whether this representational enhancement extends to encoding the temporal order of items within sequences. Musical context significantly enhanced sequential learning behaviors across multiple measures. During encoding, participants showed consistently higher retrieval accuracy for music-accompanied sequences, particularly in later encoding runs (**Figure 6c**). This enhanced online learning translated to robust subsequent sequential memory retrieval: participants reconstructed music-accompanied sequences more accurately than the control sequences in the final retrieval task (**Figure 7a**). These behavioral improvements suggested that musical temporal structure facilitated encoding of temporal order relationships, motivating our investigation of the underlying hippocampal mechanisms.

Analysis of how hippocampal representations related to enhanced behavioral performance revealed a significant mechanistic transformation (**Figure 6a-b**). Pattern similarity between adjacent items showed opposite relationships with subsequent retrieval performance across conditions. This reversal reveals fundamentally different “failure modes” for memory formation with and without musical context, illuminating how external structure transforms hippocampal computation.

In the control condition when sequences lacked additional temporal context cues, forgotten items were characterized by excessive pattern similarity with subsequent items, likely reflecting failures in pattern separation where the hippocampus couldn’t adequately distinguish between sequential items, leading to overgeneralization during encoding, aligning with its role in distinguishing similar items or items that took place at similar timing or location (Brown et al., 2014; Ranganath & Hsieh, 2016; Yassa & Stark, 2011).

Conversely, in the music condition, forgotten items showed weaker representational similarity with subsequent items than remembered items. This suggested that forgetting in the music condition appears to reflect insufficient contextual binding—items that failed to be strongly linked to their shared musical context may be ‘isolated’ in memory (Howard & Kahana, 2002; Polyn et al., 2009). This pattern of results aligns with established findings in context-dependent memory research, where degradation of contextual information significantly impairs retrieval processes (Smith & Vela, 2001). Items that mapped to music’s added contextual structure were more likely to be remembered in sequence.

Critically, music did not just help segment sequences: hippocampal patterns in the music condition contained more reliable information about item position within the sequences as revealed by multivariate classification analysis (**Figure 7.b**). This enhanced positional coding suggests that the consistent temporal framwork provided by familiar melodies does more than simply mark boundaries – it scaffolds encoding of temporal relationships between items. The temporal predictability of familiar music may provide an external ‘reference frame’ that supports more precise encoding of item positions.

One potential concern was whether enhanced positional encoding and increased within-sequence similarity (**Figure 5**, top) might represent contradictory findings. However, these effects reflect very different aspects of hippocampal representations captured by different analytical approaches. RSA measures global pattern similarity across all voxels, reflecting the overall signal structure, while MVPA identifies the specific dimensions, potentially a small subset of features, that optimally discriminate between positions. Musical context may add a common temporal template that increases overall similarity (like adding a carrier signal) while preserving or enhancing the discriminative features that encode ordinal information (like maintaining distinct phase relationships).

The hippocampus has long been recognized as critical for temporal order memory, with extensive evidence for neural populations that encode sequential position information (Hsieh et al., 2014; MacDonald et al., 2011). Time cells in the hippocampus fire at specific temporal delays, creating a neural timeline that supports sequence learning (Eichenbaum, 2014). Indeed, by representing temporal context through gradually changing patterns of activity, the hippocampus may enable discrimination between events that occur at different times (Howard & Eichenbaum, 2015; Mankin et al., 2012). The enhanced positional classification accuracy we observed extend this literature by suggesting that temporal structure signals from one modality (music) provide additional scaffolding for these context representations, creating more distinct and stable temporal codes in the other modality (visual).

This cross-modal enhancement connects to emerging work on how different sensory modalities interact during sequence learning. Studies have shown that motor temporal patterns can influence visual sequence processing (Gasser & Davachi, 2023), and that the hippocampus processes and integrates multisensory order information (Radvansky et al., 2021); Previous cross-modal studies have revealed important principles of temporal synchrony and binding (Shams & Seitz, 2008), and our findings extend this work by demonstrating that temporally predictable auditory context can scaffold visual sequential memory through hippocampal mechanisms. Furthermore, building on established single-modality paradigms where participants learn statistical relationships (Schapiro et al., 2013; Turk-Browne et al., 2009), we demonstrate that cross-modal temporal structure can also enhance probabilistic learning from noisy input. Complementing prior work examining specific aspects of hippocampal computation (Schapiro et al., 2016), we demonstrate how external scaffolding can simultaneously optimize both binding and separation mechanisms, extending beyond simple multisensory binding to facilitate hierarchical organization of memory in another modality.

### Broader Implications

The coordinated enhancements of boundary detection, representational optimization, and positional encoding demonstrate that external predictable temporal structure from musical context can reorganize multiple aspects of hippocampal computation. This finding has broad implications for understanding cross-modal learning and memory organization, suggesting that appropriate temporal context can provide a framework for structuring new memories. The fact this process can occur without explicit instruction to connect modalities suggests our study taps fundamental computational principles whereby the brain leverages available temporal structure to optimize memory organization across sensory domains.

Our findings also provide mechanistic insight for clinical and educational applications. Musical interventions have demonstrated benefits in Alzheimer’s disease and mild cognitive impairment (Moussard et al., 2012; Särkämö et al., 2008; Simmons-Stern et al., 2010). Our results help explain these effects: early Alzheimer’s disease preferentially affects CA1 and entorhinal cortex while relatively sparing CA3 and dentate gyrus in initial stages (Braak & Braak, 1991; Small et al., 2011). The enhanced within-sequence binding we observed in CA3 and the improved pattern separation in dentate gyrus suggest that musical scaffolding could recruit these preserved circuits to compensate for early CA1 dysfunction. Similarly, in educational contexts musical training enhances language acquisition (Patel, 2011), effects that align with our demonstration of cross-modal temporal scaffolding. Our finding that musical context enhanced both item binding and sequential position encoding provides mechanistic insight into why musical training improves statistical learning of linguistic patterns (François et al., 2013; Francois & Schön, 2011).

### Why Music?

Why does familiar music provide such robust enhancement for sequence learning? We argue that the answer lies not in music’s auditory properties per se, but in its unique temporal predictability that creates a coherent organizational template.

Unlike monotonic tones or random sounds or random colors, familiar music contains predictable hierarchical structure at multiple temporal scales, from rhythmic patterns to melodic phrases to overall song structure (Koelsch et al., 2019). This provides both local temporal cues (enabling boundary detection) and global organizational frameworks (supporting positional encoding). Our previous work directly demonstrates this distinction: familiar music enhanced temporal order memory, while temporally matched monotonic tones provided no benefits, confirming that structured temporal predictability, not mere auditory stimulation, drives these effects (Ren et al., 2024; Ren & Brown, 2025). Those studies demonstrated the behavioral effectiveness in the present report likely depend on having a known temporal template that can organize new learning, consistent with schema theory’s prediction that prior knowledge structures facilitate new learning (van Kesteren et al., 2012). This suggests a broader principle: any structured, familiar temporal pattern could enhance sequence learning. Indeed, existing motor sequence knowledge enhances episodic memory formation (Gasser & Davachi, 2023), and regular rhythmic presentation can improve item recognition memory (Jones & Ward, 2019); we thus postulate that our present findings should extend beyond Western music to when other similarly-regular and schematic temporal structures are present during learning.

A potential limitation is our use of silence rather than an active auditory control condition. However, this design choice was informed by our previous works which already demonstrated that familiar music produces greater behavioral and hippocampal enhancement compared to monotonic tone controls (Ren et al., 2024; Ren & Brown, 2025), indicating that benefits arise from structured temporal patterns rather than mere rhythmic auditory stimulation. Moreover, silence represents the natural baseline for most statistical learning paradigms and enables direct comparison with established sequence learning literature. Several lines of evidence support that our effects reflect cross-modal temporal scaffolding rather than simple auditory responses: (1) hippocampal enhancement occurred in a region not typically engaged during familiar music processing alone (Ren & Brown, 2023), (2) effects emerged despite explicit instructions to focus solely on visual sequences, suggesting automatic cross-modal transfer rather than divided attention, and (3) the specific neural patterns observed (enhanced boundary detection, subfield-specific representational changes, and reversed pattern-memory relationships) are consistent with hierarchical temporal scaffolding mechanisms rather than general auditory arousal. Finally, our MSM model shows how the empirical patterns observed can be reproduced by modeling temporal structure parameters rather than auditory-specific mechanisms, further supporting cross-modal scaffolding as the underlying principle. Nevertheless, our findings highlight clear opportunities for future studies to attempt to isolate the contributions of temporal predictability versus familiarity using musical and non-musical stimuli.

An interesting related consideration is the potential influence of participants’ attention towards music. While participants were instructed to focus solely on visual sequences, music may have captured some attention despite these instructions. However, several aspects of our findings argue against simple attentional explanations: (1) if music merely served as a distractor, it should impair rather than enhance visual sequence learning, (2) the specific neural patterns we observed (enhanced hippocampal binding, improved positional coding) reflect functional changes consistent with scaffolding rather than divided attention, and (3) the anatomical specificity across hippocampal subfields suggests systematic enhancement of memory processes rather than general arousal or attention effects.

## Conclusion

This study reveals fundamental mechanisms by which temporal structure from a stimulus stream can reorganize memory formation in another. By showing that familiar music simultaneously enhances hippocampal boundary detection, optimizes the balance between pattern separation and completion across anatomically distinct subfields, and provides cross-modal scaffolding for sequential position encoding, we demonstrate evidence for computational principles whereby environmental temporal structure can coordinate hippocampal memory circuit dynamics to enhance learning beyond what is achievable through single-modality approaches. Musical context created conditions where contextual similarity became a stronger predictor of memory success, demonstrating that external scaffolding doesn’t simply boost memory strength but reorganizes how neural representations support cognition. These findings advance our understanding of memory organization to encompass how intrinsic hippocampal computations are influenced by our sensory environment structure, with immediate implications for educational interventions and precision approaches to cognitive rehabilitation. More broadly, this work with music highlights mechanistic relationships that may be general principles through which structured environmental cues in one modality can optimize human cognitive function and enhance memory formation in another.

